# Non-REM parasomnia experiences share EEG correlates with dreams

**DOI:** 10.1101/2023.11.10.565325

**Authors:** Jacinthe Cataldi, Aurélie M. Stephan, José Haba-Rubio, Francesca Siclari

**Affiliations:** Center for Investigation and Research on Sleep, Lausanne University Hospital, Switzerland; The Sense Innovation and Research Center, Lausanne and Sion, Switzerland; The Netherlands Institute for Neuroscience, Amsterdam, The Netherlands

**Keywords:** disorders of arousal, parasomnia, sleepwalking, sleep terror, confusional arousals, dream, awakening, arousal, slow wave sleep, consciousness, amnesia, high-density EEG

## Abstract

Sleepwalking and related parasomnias result from sudden and incomplete awakenings out of slow wave sleep. Clinical observations suggest that behavioral episodes can occur without consciousness and recollection, or in relation to dream-like experiences. To understand what accounts for these differences in consciousness and amnesia, we recorded parasomnia episodes with high-density EEG and interviewed participants immediately afterwards. Compared to reports of unconsciousness (19%), reports of conscious experience (81%) were preceded, during prior sleep, by high-amplitude slow waves in anterior cortical regions and an activation of posterior cortical regions. Reduced posterior slow wave activity was also present during the episode when patients displayed elaborate behaviours in relation to dream-like scenarios. Amnesia for the experience (25%) was modulated by right medial temporal activation during prior sleep and fronto-parietal slow wave activity during the episode. Thus, the neural correlates of parasomnia experiences are similar to those previously reported for dreams and therefore likely reflect core physiological processes involved in sleep consciousness.

## Introduction

Sleepwalking (somnambulism) and related parasomnias are enigmatic conditions characterized by sudden but partial awakenings out of Non-rapid eye movement (NREM) sleep^1^, during which affected individuals may interact with their environment in an altered state of consciousness^2–4^. Behaviors during parasomnia episodes can be short and simple, such as sitting up and talking, or more complex, like sustaining a conversation, leaving the bed, or manipulating objects. In extreme cases, sleepwalkers have been reported to drive, commit sexual assault, homicide and pseudo-suicide, resulting in personal tragedies and legal dilemmas^5–7^. Although rare, these instances invariably raise the question of whether sleepwalkers are conscious, and if yes, what they experience during parasomnia episodes.

Anecdotal reports describing ‘dream-like’ experiences during sleepwalking date back to the 18^th^ century^5^, but were largely dismissed in the 1960s, when the first electroencephalographic (EEG) recordings of somnambulistic episodes showed that this condition occurred out of slow wave sleep^8,9^, a sleep stage that at the time was considered largely dreamless and in stark opposition to the recently discovered REM sleep^10,11^. Although it was later acknowledged that patients could report ‘apparent dream recall’ after parasomnia episodes^12,13^, some questioned that this memory truly reflected dreaming^9^, and the view that sleepwalking predominantly represented a largely unconscious, ‘ambulatory automatism’^8^ prevailed. In subsequent years, paralleling the growing evidence for NREM dreaming^14^, dream-like experiences during NREM parasomnia episodes were increasingly reported^3,15,24–26,16–23^. These experiences most often consisted in threatening scenarios^15–17,27^ and could encompass hallucinations^20–22^, delusions^23,24^ and behaviors in line with the ‘dream’ content. Larger studies, in which patients were asked to describe, over a lifetime, what they had experienced during NREM parasomnia episodes revealed that 66-94% of adults could report at least one experience^19,22,27^, while this rate was only about a third in children^25^. There is also anecdotal evidence of NREM parasomnia experiences with minimal or partial consciousness, and with variable degrees of amnesia^6^. However, almost all of these studies relied on retrospective recall of experiences over a lifetime, and are therefore prone to recall biases, especially since it is well known that dream-related memories vanish rapidly^28^. In one of the rare studies in which conscious experiences were assessed immediately after parasomnia episodes in patients with sleep terrors, vivid experiences were reported in 58% of cases, and vague recall in 7%^12^. Thus, the level of consciousness and amnesia associated with NREM parasomnia episodes is likely variable than previously thought, ranging from episodes with no or minimal consciousness or recall, to vivid, dream-like experiences.

Our first aim was therefore to systematically assess and quantify this variability by interviewing patients immediately after NREM parasomnia episodes about their experiences. Our second aim was to understand which brain activity changes, at the cortical level, could account for the variability in consciousness and amnesia associated with parasomnia episodes. Previous studies using nuclear imaging and intracranial EEG recordings documented patterns of wake- and sleep-like activity in different brain regions during NREM parasomnia episodes^4,29–32^, but these techniques are not readily available to study a large number of episodes. It remains therefore unclear how these dissociated brain activity patterns relate to conscious experience. Here we took advantage of high-density (hd-) EEG recordings to record parasomnia episodes, which we combined with a serial interview paradigm, a method that has previously allowed us, in healthy sleepers, to document brain activity patterns that distinguish unconsciousness from dreaming in both REM and NREM sleep^33,34^. Two previous case reports have used high-density EEG to image brain activity before and during parasomnia episodes^35,36^, demonstrating feasibility of the approach. More specifically, our previous studies showed that compared to reports of unconsciousness, reports of dreaming were preceded by a regional EEG activation in parieto-occipital brain areas (grouped under the name ‘posterior hot zone’)^33^, and in NREM sleep, by high-amplitude frontal slow waves (K-complexes/type I slow waves) that are likely related to arousal systems^34^. We hypothesized that if these EEG features reflect core physiological processes involved in sleep consciousness, they should also distinguish parasomnia episodes with and without conscious experience.

## Results

### Patients

Twenty-two patients with disorders of arousal were included [14 females, aged 26.9 ± 5.3 yrs (average ± SD), range 18.3-36.3 yrs, see supplementary material Text S1 for additional information on patients]. Twenty of these participants were included in a previous publication ^37^ while two participants were newly recruited. Participants underwent two high-density EEG sleep recordings: a first nighttime recording and a second daytime recording after 25h of sleep deprivation, during which acoustic stimuli were administered to increase the complexity and incidence of parasomnia episodes ^38–40^ (see methods section for details). Immediately after each parasomnia episode, the examiner called the patients by name and asked about their most recent experience (based on^33,41^). When participants did not report a conscious experience (CE), they had to clarify whether they had experienced something but could not recall/report the content of the experience (conscious experience without report of content, CEWR), or whether they had not experienced anything (no experience, NE).

### Parasomnia episodes

102 potential parasomnia episodes [19 patients, 5.3 ± 3.54 episodes per patient, range 1-12] were recorded and examined by three independent raters. Of these, 75 (73.5%, 18 patients) were unanimously rated as parasomnia episodes by all three experts and were included in the analyses. Parasomnia episodes mostly consisted of short confusional arousals. None of the patients left the bed, although the setup allowed them to do so. Episodes lasted on average 28.6s (+/-18.8s, median 22.0s, range 3 -108s), and occurred 149.7 (+/-91.5) min after lights off (range 8–373 min). Of the 75 episodes unanimously rated as parasomnia episodes, 48 (64.0%) occurred spontaneously and 27 (36.0%) were provoked by sounds. 37 episodes (49.3%) occurred during the baseline night and 38 (50.7%) during the recovery night. 69 episodes (92.0%) occurred out of stage N3, and 6 (8.0%) out of stage N2.

### Behavioral features of parasomnia episodes

Most episodes had a sudden onset and were characterized by eye opening and manifestations of surprise, including orienting behaviour (exploratory head and eye movements) and expressions of perplexity, as well as somniloquia (Table 1). Interestingly, manifestations of surprise were not only observed at the onset of provoked parasomnia episodes, but also with a similar frequency in spontaneous episodes, when no sound was played, consistent with our previous observation that these two types of episodes are behaviorally indistinguishable^37^. Indeed, a generalized linear model was not able to predict the provoked vs. spontaneous nature of the behaviour based on the presence of surprise features (included as fixed factor while accounting for subject identity as random factor, [(*χ*^2^(1) = 0.22, p = 0.64), see methods and Table S1 for information on statistical models].

More complex behaviours were seen in longer episodes (> 30s) and included searching (going through the bed sheets, looking under the bed), gesticulating (trying to catch or grasp non-existent things, military salutation), sustaining imaginary conversations (i.e., alternatively speaking and pausing, as if listening), visually scanning the environment for continued periods and/or pointing towards (imaginary) objects. Apparent hallucinations and/or perceptual illusions were present in 52% of episodes.

**Table 1:** Behavioural features of parasomnia episodes. Presence/absence of feature (%) refers to instances in which at least two of the three raters agreed.

| Feature | Presence<br>N (%) | Absence<br>N (%) | No agreement<br>N (%) | Other<br>reason (n) |
| --- | --- | --- | --- | --- |
| Eye opening | 60 (80%) | 4 (5%) | 2 (3%) | not visible (4), both open/closed (9) |
| Sudden onset | 49 (65%) | 24 (32%) | 2 (3%) | - |
| Perplexity | 48 (64%) | 21 (28%) | 2 (3%) | not visible (4) |
| Orienting behaviour | 45 (60%) | 28 (37%) | 1 (1%) | not visible (1) |
| Somniloquia | 43 (57%) | 32 (43%) | 0 | - |
| Interactions with environment | 29 (39%) | 42 (56%) | 4 (5%) | - |
| Startle-like reaction at onset | 28 (37%) | 45 (60%) | 2 (3%) | - |
| Expression of fear | 17 (23%) | 53 (71%) | 2 (3%) | not visible (3) |
| Smiling or laughing | 3 (4%) | 66 (88%) | 2 (3%) | not visible (4) |

### Consciousness and parasomnia episodes

Of the 75 unanimously scored episodes, 57 (76%) were followed by an interview about conscious experiences. Instances without interview occurred either because the experimenter failed to recognize the parasomnia episode as such during the recording, or because the patient fell asleep too quickly after the episode to answer questions. CE were reported in 56.1 % (n=32, 14 patients), CEWR in 24.6% (n=14, 8 patients) and NE in 19.3% (n=11, 7 patients Fig. 1A and B).

Reported conscious experiences included constructed delirious scenarios (n=25), in which the patients were trying to prevent impending danger or its consequences (like needing to save ladybugs from dying, having to find one’s baby daughter who had fallen off the bed, struggling because being enclosed in bedsheets, impression that a piece of furniture was coming down from the ceiling), but also more ordinary scenarios (telling the experimenter how to fall asleep without making any sound, needing to go to an appointment with a therapist, seeing one’s daughter vomit), as well as isolated imagery (a shower, green pastures, cookies, landscapes, n=5) or thoughts (about taxes, bills, n=2). Only 2 out of 18 CE reports following provoked episodes (11%) contained a possible reference to the alarm sound: in one case the patient mentioned a conversation during which she hear a sound, leading her to turn around rapidly (which she did not do in reality), and in the other case the patient mentioned “a noise, like a sound”, and being scared of it, she also saw the image of an alarm. Interestingly, in some spontaneous parasomnia episodes, patients also acted as if they had perceived a sudden sensory stimulus, for instance by asking a question like ‘What’? ‘Hmm’? or ‘What did you say?’. A clear correspondence between the reported experience and the behaviour was evident in 47% cases (n=15, coherent CE, Video S1), in 13% (n=4) it was only partially apparent, in 34% (n = 11) a correspondence was not apparent, and in 6% (n = 2) the report was totally incompatible with the behaviour (the last two categories were grouped in the incoherent CE category, for an example see Video S2). A table with reports of experiences and associated behaviours is provided in Table S2.

**Fig 1.**
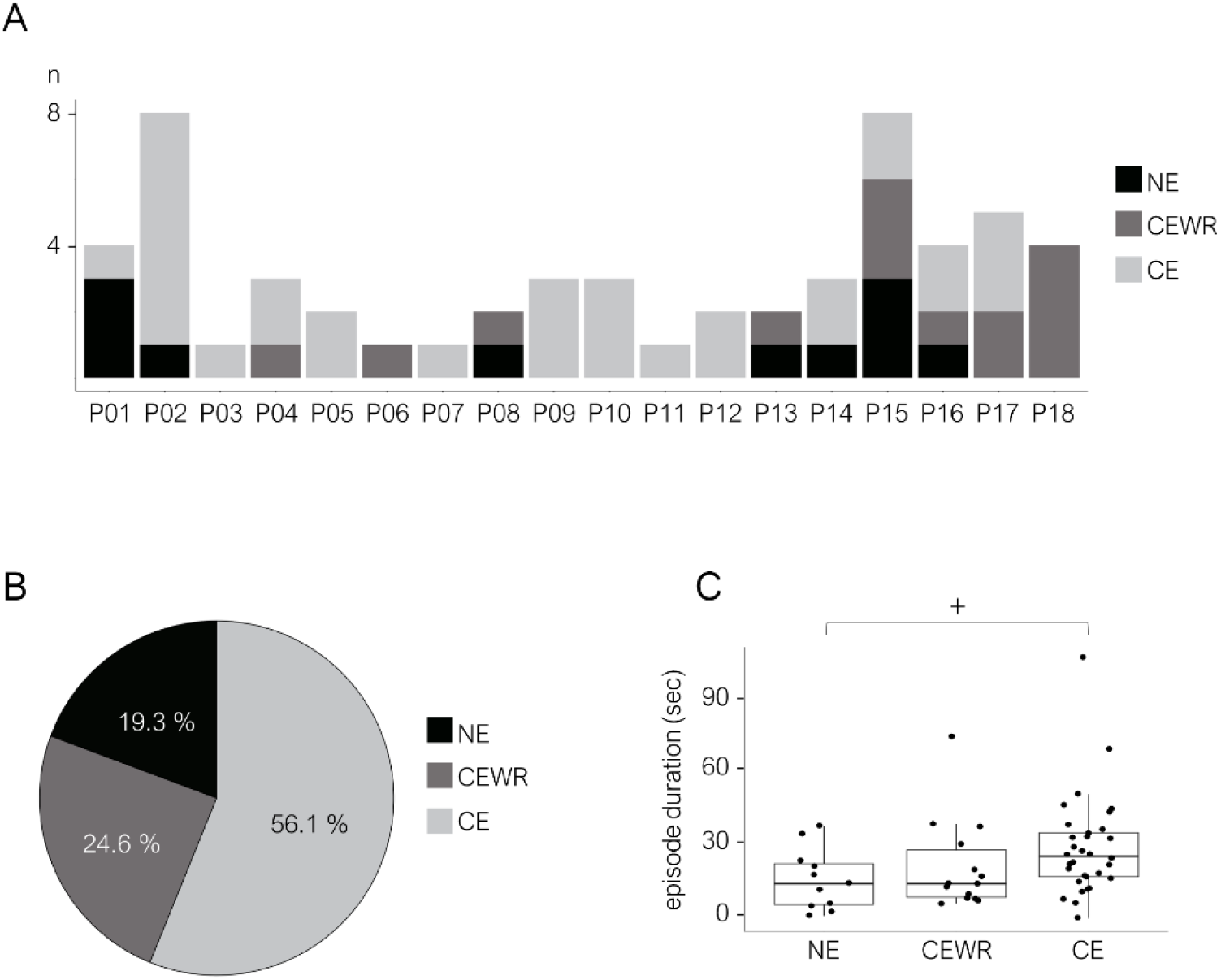
Consciousness and NREM parasomnia episodes. **A.** Absolute number of parasomnia episodes per patient with report of conscious experience (CE), conscious experience without recall (CEWR) or no experience (NE). **B.** Relative frequency (%) of CE, CEWR and NE across all trials. **C.** Duration of episodes with CE, CEWR and NE. Each black point represents a parasomnia episode, boxes display the 25^th^ to 75^th^ percentile of data, horizontal bars indicate the median, vertical bars the confidence interval. +: p<0.1.

Episodes after which patients reported no experience (N=11, Video S3) were all but one characterized by at least one manifestation of surprise (rapid onset, startle-like movement sequence, orienting behaviour, and/or perplexity), whether a sound had been played or not, and were sometimes accompanied by somniloquia (N=9), a fearful facial expression (N=4) and/or repetitive hand movements (N=2). When asked, most patients said that they were not aware of having displayed any behaviour at all (n=5), while one patient reported ‘simply’ having turned in bed, although he had sat up in bed with a frightened facial expression. The category of report (CE vs NE or CE vs CEWR) did not vary as a function of the provoked vs. spontaneous nature of the episode or its occurrence during the first vs. second recording, the time since lights off or the time since the first sleep onset (Table S1). Median durations of CE episodes were longer than CEWR and NE episodes, (see Table S3 but these contrasts did not reach statistical significance (CE vs. CEWR: p = 0.546, z = 0.603 and CE vs. NE: p = 0.067, z = 1.832, Fig. 1C). Episodes for which a clear correspondence between the report and the behaviour was observed were significantly longer than NE (18.8 +/-12.15 secs vs 37.13 +/-22.17 secs, duration: *χ*^2^(1) = 2.24, p = 0.024).

### EEG correlates of parasomnia experiences

Compared to parasomnia episodes with NE, those with CE were predicted by lower delta (1-4 Hz) and higher beta power (26-34 Hz) in posterior cortical regions in the 20s of sleep preceding movement onset (Fig. 2 A and B, left, see Fig. S1 for results at the scalp level). These differences were centered on primary visual cortices, extended inferiorly to occipital-temporal areas, anteriorly to medial temporal regions and superiorly to include parts of the precuneus and the posterior cingulate cortex, similar to the EEG correlates previously reported for NREM and REM sleep dreams using the same methodology^33^ (see overlap maps with respect to previous study in supplementary Fig. S2 and Table S4 for cluster statistics). Differences in beta power additionally included lateral temporal areas and parts of the pre-and postcentral gyri. Parasomnia episodes with CE were also predicted, compared to those with NE, by lower delta power in posterior cortical regions *during* the episode (+4s to +20s after movement onset, Fig. 2A right, bottom panel, see methods section for rationale behind choice of timeframes); however, compared to the previous contrast these differences were mainly left sided and comprised mostly lateral parietal and temporal areas (angular and supramarginal gyrus, inferior parietal lobule, paracentral lobule, precuneus), as well as parts of the postcentral gyrus on the right side. Very similar findings to the CE-NE contrast were obtained when comparing instances in which patients reported experiences, but did not remember the content (CEWR), with NE, although results were only present in the sleep EEG and were statistically weaker because fewer episodes were available for comparison (Fig. S3). Thus, decreased delta power in parieto-occipital cortices precedes parasomnia episodes with CE compared to NE, irrespective of whether the content is later remembered.

Compared to parasomnia episodes with NE, those with CE were also predicted by anterior and central slow waves with a higher amplitude, a steeper slope and fewer negative peaks, and by fewer slow waves in posterior brain regions (Fig. 3 and S4), a slow wave constellation that also precedes reports of dreaming^12^. Similar to previous results for delta power in this study, CE were also associated with smaller, shallower and shorter slow waves in left posterior regions *during* parasomnia compared to NE.

**Fig 2.**
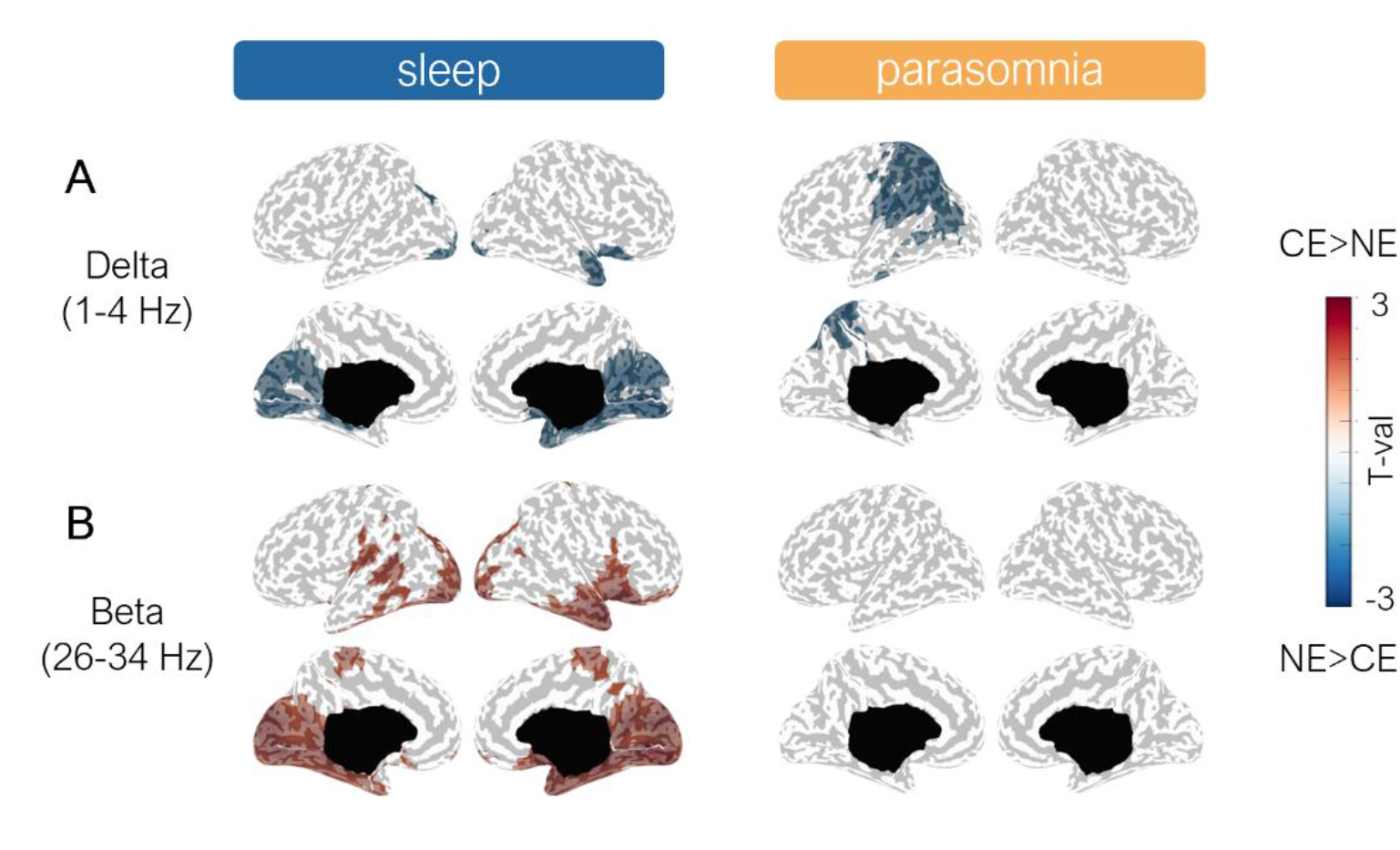
Conscious experience (CE) vs. no experience (NE): spectral power. **A.** Cortical distribution of statistical differences in delta (**A**) and beta (**B**) power (t-values, Wald statistics) for CE vs. NE. Right column: 20 seconds of sleep preceding movement onset (CE: n=32, NE: n=11). Left column: from 4 to 20 seconds after movement onset, corresponding to the parasomnia episode (CE: n=31, NE: n=10). Only voxels with significant effects appear colored. LL, left lateral; RL, right lateral; LM, left medial; RM, right medial.

**Fig. 3:**
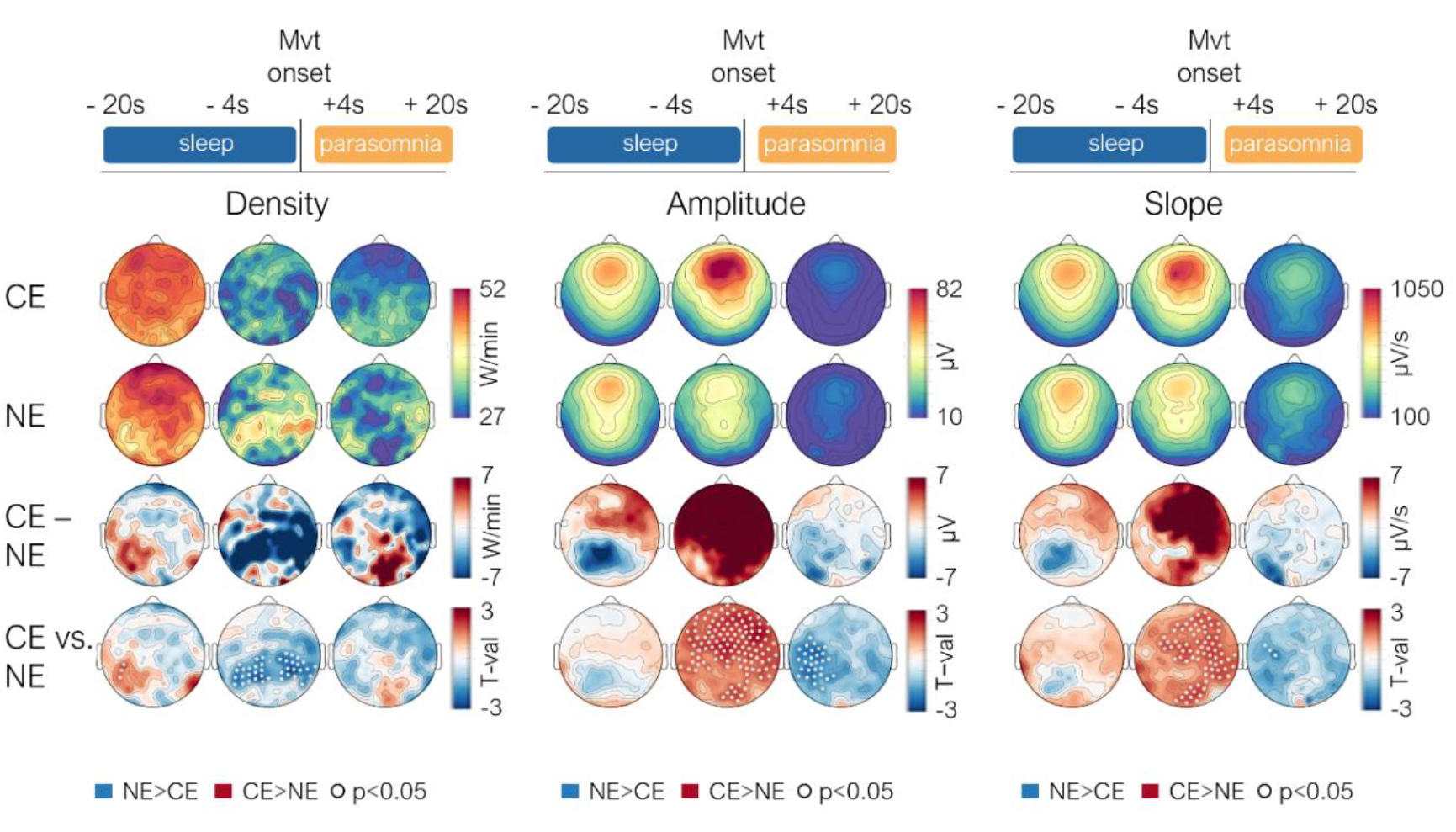
Conscious experience (CE) vs. no experience (NE): slow wave parameters. Topographical distribution of slow wave parameters averaged across subjects for CE (N=14 subjects, 32 episodes) and NE (n=7, 11 episodes), of their absolute difference (CE minus NE), and of t-values (Wald statistics, CE vs. NE) for three timeframes (sleep from -20 to -4s before movement onset (n= 43 trials), sleep from -4s to movement onset (n= 43 trials) and the parasomnia episode from +4s to +20s (n= 41 trials)). 175 innermost channels are displayed. Density=number of slow waves per minute, Amplitude= absolute amplitude of maximum negative slow wave peak, Slope = slope of negative to positive deflection of slow wave. Mvt onset= movement onset.

### EEG correlates of parasomnia experiences that were coherent and incoherent with behaviour

We then asked whether the correlates of parasomnia experiences that were coherent with the observed behaviour (CCE) were different from those for which no coherence was observed (ICE). Compared to NE, both coherent and incoherent episodes displayed differences in posterior cortical regions in the scalp sleep EEG that were also seen when contrasting all experiences with NE, consisting in either increased beta power or reduced slow wave amplitude/slope in posterior regions (Fig. 4 and S5), although these differences were statistically weaker because fewer trials were considered, and no clear results emerged from source reconstruction maps. However, only behavioural episodes which were coherent with the reported experiences showed higher amplitude (type I) slow waves in frontal-central regions immediately prior to movement onset and lower delta power and slow wave amplitudes/slopes *during* parasomnia episodes compared to NE. Thus, the previously documented differences in delta power between CE and NE *during* parasomnia episodes were likely driven by those episodes for which the behaviour was coherent with the report, while the ones for which no coherence was observed were associated with differences well before movement onset.

**Fig. 4:**
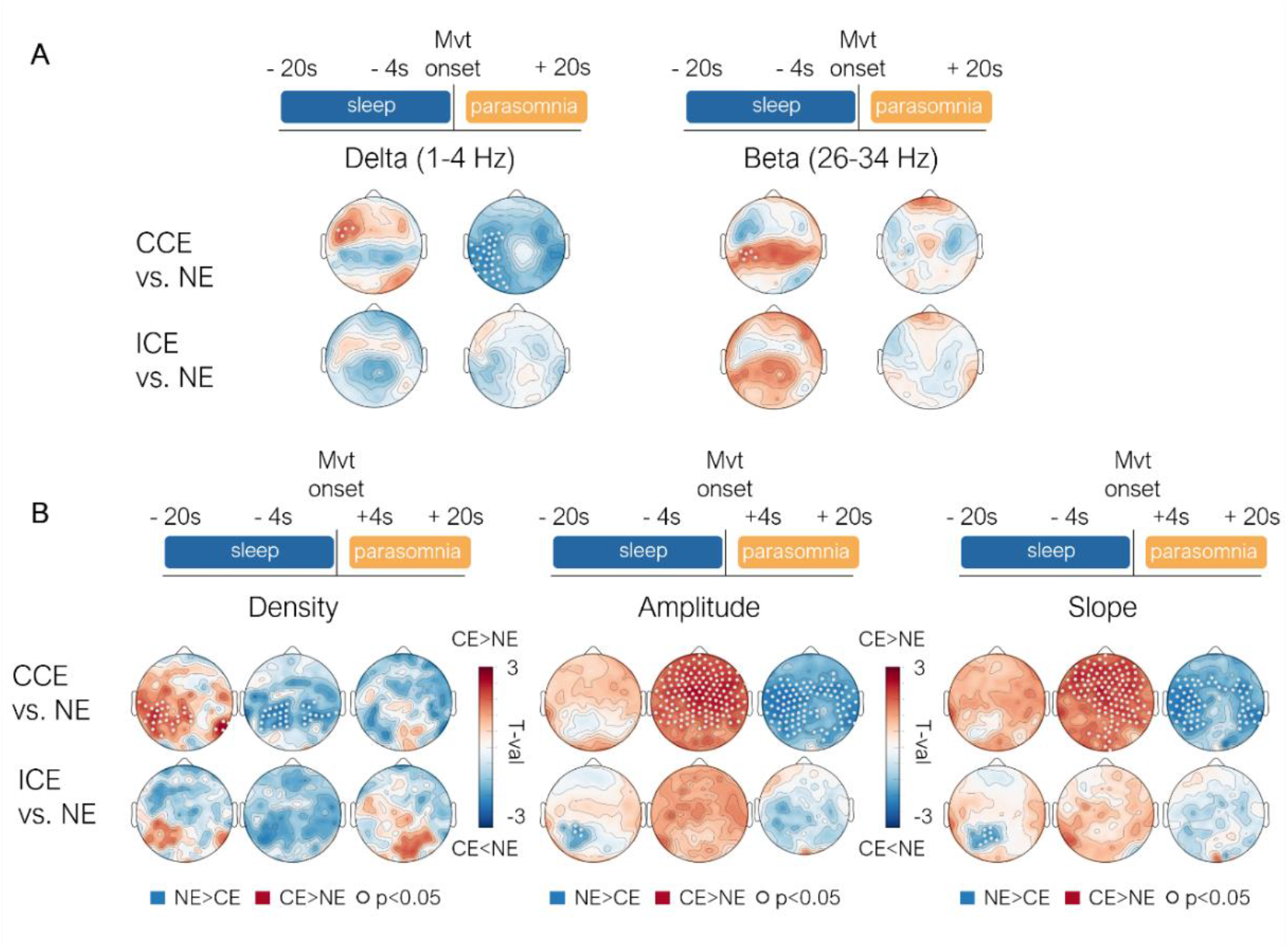
Coherent (CCE) and incoherent (ICE) conscious experiences vs. no experience (NE). **A.** Topographical distribution of statistical differences in delta and beta power (t-values, Wald statistics) for CCE vs. NE (top row in each panel; n=15 vs. 11 episodes) and ICE vs. NE (bottom row in each panel; n=13 vs 11), shown for the EEG corresponding to sleep (left) and parasomnia episode (right). **B.** Topographical distribution of statistical differences in slow wave parameters (t-values, Wald statistics) for CCE vs. NE (top row in each panel; n=15 vs. 11 episodes) and ICE vs. NE (bottom row in each panel; n=13 vs 11 episodes) shown for three timeframes (sleep from -20 to -4s before movement onset, sleep from -4s to movement onset and parasomnia episode from +4s to +20s).175 innermost channels are displayed. Density=number of slow waves per minute, Amplitude= absolute amplitude of maximum negative slow wave peak, Slope =slope of negative to positive deflection of slow wave. Mvt onset= movement onset.

### EEG correlates of recall of parasomnia experiences

Next, we tried to determine the correlates of the recall of the experience by comparing instances of CEWR and CE. Compared to CEWR, CE with recall were preceded, in the pre-episode sleep, by lower delta power and higher beta power in a circumscribed anterior region of the right medial temporal lobe, estimated to comprise the hippocampus, parahippocampal gyrus and amygdala (Fig. 5 A and B left). CE with recall were also predicted by lower delta power in a relatively widespread fronto-parietal area *during* parasomnia episodes compared to CEWR (Fig. 5A right, see Fig. S6 for results at the scalp level), comprising medially the precuneus, posterior and central cingulate gyrus, the paracentral lobule and the superior frontal gyrus, extending laterally on the right hemisphere to the pre-and postcentral gyri, middle frontal gyri and the insula. The analysis of slow wave parameters confirmed that recall of the content of the experience was mostly predicted by widespread slow wave differences after movement onset during the episode (Fig. 6 and S7), while no such differences were seen in the sleep EEG.

**Fig. 5:**
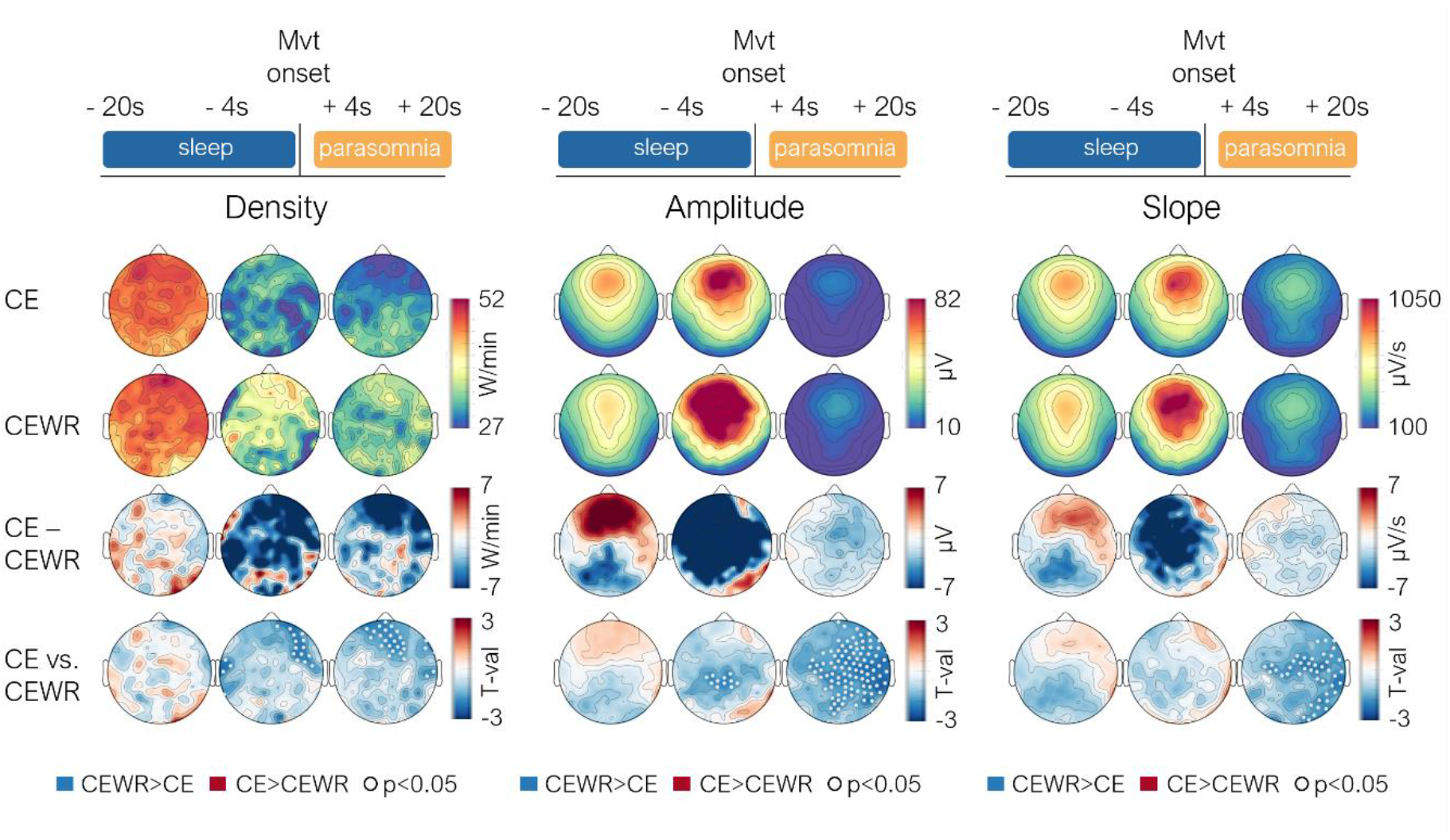
Conscious experience (CE) vs. conscious experience without recall of content (CEWR): spectral power. Cortical distribution of statistical differences in delta (**A**) and beta (**B**) power (t-values, Wald statistics) for CE vs. CEWR. Right column: 20 seconds of sleep preceding movement onset (CE: n=32, CEWR: n=17). Left column: from 4 to 20 seconds after the movement onset, corresponding to the parasomnia episode (CE: n=31, CEWR: n=17). Only voxels with significant effect appear colored. LL, left lateral; RL, right lateral; LM, left medial; RM, right medial. Mvt onset= movement onset.

**Fig. 6:**
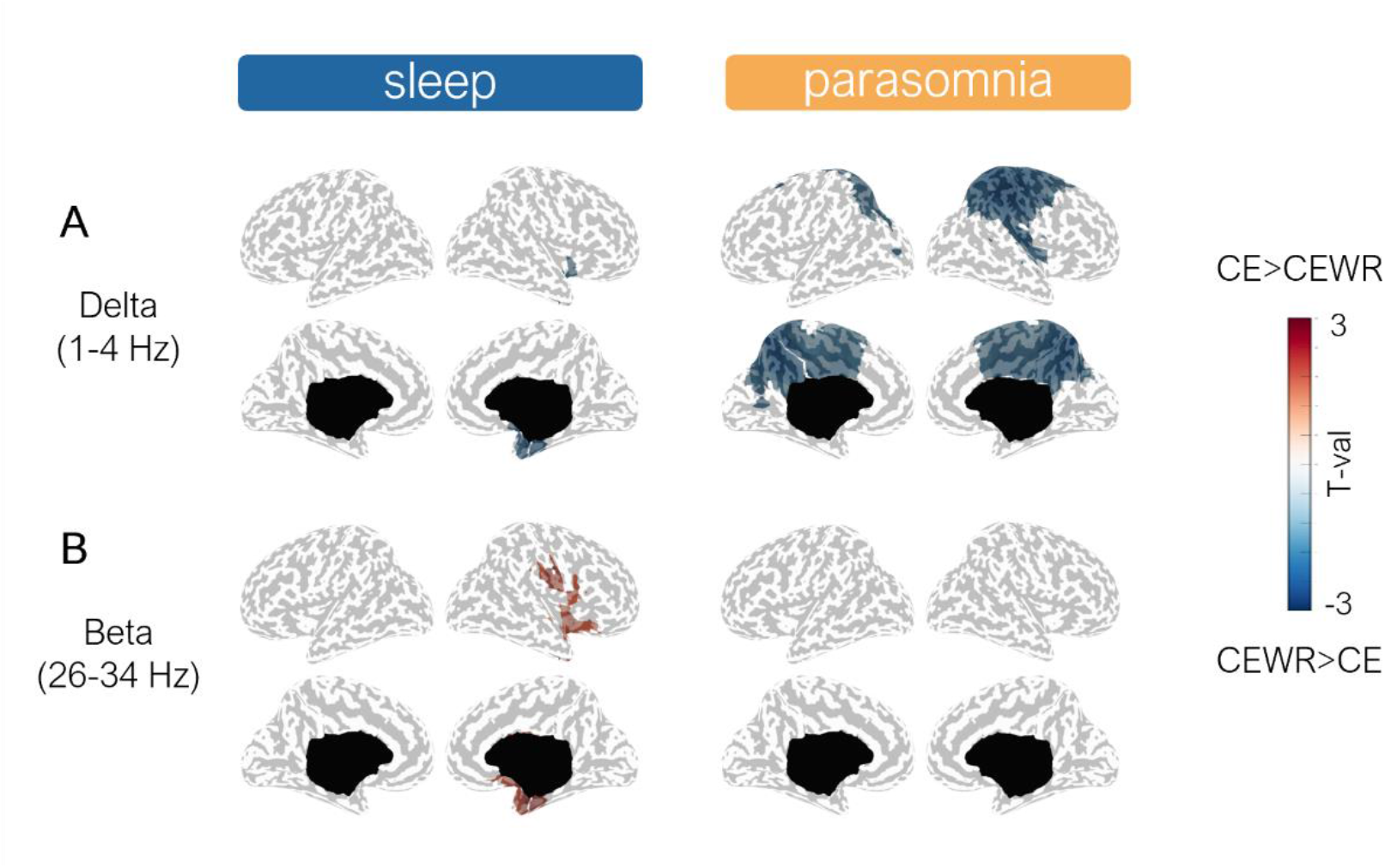
Conscious experience (CE) vs. conscious experience without recall of content (CEWR): slow wave parameters. Topographical distribution of slow wave parameters averaged across subjects for CE (N=14 subjects, 32 episodes) and CEWR (n=8, 17 episodes), of their absolute difference (CE minus CEWR), and of t-values (Wald statistics, CE vs. CEWR) for three timeframes (sleep from -20 to -4s before movement onset (49 trials), sleep from -4s to movement onset (49 trials) and the parasomnia episode from +4s to +20s (48 trials)). 175 innermost channels are displayed. Density=number of slow waves per minute, Amplitude= absolute amplitude of maximum negative slow wave peak, Slope = slope of negative to positive deflection of slow wave. Mvt onset= movement onset.

## Discussion

Our study revealed that the degree of consciousness and amnesia associated with NREM parasomnia episodes is variable, both between and within individuals. Patients’ reports ranged from no experience to dreamlike scenarios characterized by delusional thinking, multisensory hallucinations and volitional interactions with the environment. They reported a conscious experience after 81% of episodes, could clearly recall the content of the experience in 56% and reported no experience (unconsciousness) in 19%. These results are remarkably consistent with those of a previous study documenting immediate recall of experience with clear content in 58% of NREM parasomnia episodes^12^, and more broadly with the rate of dreaming reported after awakening from NREM sleep stages 2/3 when using the same methodology^41^.

Most parasomnia episodes started abruptly, and were characterized, at their onset, by facial expressions of surprise and perplexity, as well as exploratory behaviours (eye opening, orienting eye and head movements). Intriguingly, these behaviours at onset occurred not only in response to loud sounds, but also spontaneously, and perhaps even more surprisingly, sometimes without reports of experience (consciousness). More complex and idiosyncratic behaviours emerged during longer episodes and preferentially correlated with reports of experience. Thus, the initial behaviour seen in NREM parasomnia episodes, which is similar across subjects (as also reported in^8^), likely reflects a common stereotyped activation of arousal systems, which can either occur spontaneously, or as a response to sudden arousing stimuli, and be dissociated from consciousness. The fact that almost identical behaviours, EEG patterns^37^ and mental contents are observed between spontaneous and provoked parasomnia episodes suggests that provoked episodes mimic a naturally occurring arousal process^13^. Indeed, recurrent activations of arousal systems are now known to be an integral part of NREM sleep^42,43^, as recently confirmed by studies demonstrating periodic activations of the locus coeruleus in slow wave sleep^44,45^. It is therefore not too far-fetched to assume that spontaneous, naturally occurring activations of arousal systems, albeit with a pathological timing^37^, intensity and/or motor coupling, could underlie the occurrence of NREM parasomnia episodes.

At the EEG level, reports of experience, compared to those of no experience, were preceded by an activation of posterior cortical regions during the sleep prior to the episode, similar to brain activity patterns that were found to distinguish dreaming from no experience in both REM and NREM sleep using the same methodology^33^. Parasomnia experiences were also preceded by large and steep slow waves in frontal and central regions, similar to the slow wave constellation that precedes reports of NREM dreaming^34^, although in our previous study, it most consistently distinguished experiences with and without recall of content. These large fronto-central slow waves are reminiscent of so-called type I slow waves/K-complexes, which are likely to be related to phasic activations of arousal systems ^46,47^ and differ in several aspects from so-called delta waves (type II slow waves) that constitute the “background EEG activity” of slow wave sleep and account for changes in the posterior hot zone of dreaming^34^. The fact that the relatively short and stereotyped behaviors without report of consciousness were not associated with these EEG changes could reflect a functional subcortico-cortical disconnection, and a predominantly subcortical generation of behavior in this case, similar to orienting and threat-related defence responses that can occur without participation of the cortex ^48–51^.

Compared to parasomnia episodes without report of experience, those with report of experience not only displayed reduced delta power in the posterior hot zone during prior sleep, but also *during* the episode, especially in cases in which the behavior matched the report of experience, as if the brain remained in ‘dream mode’ after the initial arousal in these cases. Interestingly, *while before* the episode differences between CE and NE were mainly localized to primary and secondary visual areas including the ventral visual stream (the ‘what’ pathway) *during* the episode they involved more parietal, higher-order, and multisensory associative areas, and in particular the dorsal visual stream involved in spatial vision (the ‘where’ pathway)^52^. It is not clear what underlies these differences in topography, but contrary to sleep, during parasomnia episodes, by definition, patients moved around and interacted with their environment. One tentative explanation is therefore that the more consistent involvement of multisensory areas and the dorsal visual stream during the episode reflects processes associated with spatial exploration and action.

Finally, our results indicate that amnesia for the content of the experience is favoured by relative right hippocampal deactivation during preceding sleep, which may prevent the encoding of conscious experiences (dreams) into episodic memory, as well as with persistence, after movement onset, of slow wave activity in fronto-parietal regions contributing to working memory. These latter regions are again similar to the ones we previously identified as being involved in the recall of NREM dreams^33^.

Taken together, our results suggest that NREM parasomnia experiences not only display the core features of dreams^22^, but that they are also associated with similar brain activity patterns, which are therefore likely to reflect fundamental neurophysiological mechanisms involved in sleep consciousness^53^. Our results also raise the possibility that arousal systems could contribute to dream contents. In this study we were able to induce parasomnia episodes with loud arousing sounds and observed that reported dream contents, as also described previously^8^, thematically often related to impending danger or threat, suggesting a relation between the alerting nature of the stimulus and the dream content. Interestingly, the actual alarm sound rarely figured in the reports of experience, but the notion of threat did so consistently, raising the possibility that activations of supramodal and non-specific (extra-lemniscal) systems, encoding the importance of the stimulus, rather than modality-specific (lemniscal) sensory pathways^54,55^, relaying the precise nature of the stimulus, preferentially contribute to the conscious experience during sleep. It is tempting to assume that a sudden activation of arousal systems, whether induced or occurring spontaneously, is interpreted as a danger signal by patients and secondarily contextualized, provided that the brain is in ‘dream’ or ‘conscious’ mode prior to the arousal and can thus ‘make something’ of it. Whether such a secondary enrichment of non-specific spontaneous arousal signals also underlies (NREM) dream contents remains to be elucidated.

A few study limitations deserved to be mentioned. Although this work provides one of the largest quantitative assessments of brain activity associated with NREM parasomnia episodes, for some subgroup analyses (i.e. those on coherent/incoherent experiences), the relatively small sample size does not allow us to draw firm conclusions. In addition, parasomnia episodes mainly consisted in confusional arousals and were recorded exclusively in adults; they are therefore not representative of the full repertoire of NREM parasomnias. Parasomnia episodes also had a variable duration, but we analyzed a fixed length, which may have resulted in less clear-cut findings between categories of consciousness. Although we used the same methodology as in our previous study on dreaming, to limit the effect of movement artifacts, we did not analyze the same frequency bands and excluded the first seconds of the episode, which may have resulted in some null findings, for instance in the beta frequency band (Fig. 2). The absence of beta power differences between the CE and NE condition during the episode (as opposed to in prior sleep) may also be related to the fact that during the episode patients were moving. Indeed, beta power reduction is commonly observed during movements ^56,57^. Finally, while our results are concordant with those of one of the largest studies on the EEG correlates of dreaming employing the same methodology, they need to be confirmed by other studies of the same type.

## Methods

### Patients

Patients presenting with disorders of arousal (confusional arousals, sleepwalking, and/or sleep terrors), as defined in the international classification of sleep disorders^1^, were recruited during clinical consultations at our sleep center, or by word of mouth. All patients underwent a medical evaluation by a board-certified sleep medicine physician and a clinical polysomnography (PSG, see Table S5). One patient experienced technical issues during the PSG and underwent an ambulatory polygraphy with respiratory and recording of leg movements instead. Patients who had presented at least one parasomnia episode during the last month were included. Exclusion criteria were major psychiatric or neurological comorbidities (identified during the medical consultation), medication (except birth control) and pregnancy. Participants gave written informed consent to participate in the study and to publish the videos provided in this manuscript. The study protocol was approved by the local ethical committee (commission cantonale d’éthique de la recherche sur l’être humain du canton de Vaud) .

### Experimental procedure

Patients underwent a first (baseline) hd-EEG nighttime recording between ∼11.30pm and ∼6.30am, during which they were continuously monitored via video and audio by an experimenter. The next day they were free to perform their usual activities but were not allowed to sleep. Compliance with the instructions was verified by wrist actigraphy. In the evening, they came back for a night of supervised total sleep deprivation in the laboratory. A sleep technician verified, via continuous video-and audio-recordings, that they did not fall asleep. The following morning, a second hd-EEG (recovery) sleep recording was carried out between ∼7.30am and ∼4pm. During this second recording, computerized pure sounds (1000 Hz, lasting 3s) were played at one-minute intervals after five minutes of stable slow wave sleep, with increasing intensities (50 to 90 dB, increase of 5dB between at each interval, adapted from^39^), until they provoked a parasomnia episode or a full awakening. This procedure has previously been shown to increase the frequency and complexity of NREM parasomnia episodes. During both the baseline and recovery sleep recordings, when a parasomnia episode occurred, the experimenter waited for the episode to end, called the patient’s name and then conducted a semi-structured interview (based on^33,41^) through intercom about the patient’s subjective experiences. If the patient did not react immediately, the experimenter called her/his name a second time. Patients were asked what they had experienced immediately before the examiner had called their name. When they did not report a conscious experience (CE), they had to clarify whether they had experienced something but could not recall the content of the experience (conscious experience without report of content, CEWR), or whether they had not experienced anything (no experience, NE).

### Rating of parasomnia episodes

The procedure for defining and rating of parasomnia episodes has been described previously^37^. In brief, the video and EEG of all “awakenings” out of slow wave sleep were extracted, regardless of whether they were associated with manifestations of parasomnia. The beginning of an episode/awakening was defined as movement onset on EMG or video, whichever occurred earlier, while the end was defined as movement cessation. In most cases, at the end of an episode patients paused, then sometimes started an interaction with the examiner to signal that they were fully awake (see, for instance, hand clapping in video S2), or resumed a sleeping position again. Three experts independently rated the behaviour seen on the videos (without the EEG) as either a parasomnia episode (as defined in the ICSD^1^) or a normal awakening. Only episodes considered as such unanimously by all the three raters were included in the analyses. Experts also rated the presence of the following features: interactions with the environment, somniloquia, manifestations of fear and of surprise (rapid onset, startle-like movement sequence, orienting behaviour, and perplexity), and, when a mental content was reported, whether it was coherent with the behaviour seen on the video or not. A feature was considered present when at least two of the three judges had rated it as such. The coherence between report and behavior was rated separately by the judges. In case of non-agreement, a discussion between the raters followed until agreement was reached. This method was chosen to reduce the complexity of the scoring procedure and to separate coherent and incoherent behavior without ulteriorly reducing the number of trials.

### EEG recordings and analysis

#### EEG recordings

Sleep was recorded with a hd-EEG system (256 channels, Electrical Geodesics, Inc., Eugene, Oregon) and a 500Hz sampling rate, in a certified sleep center, guaranteeing conditions that are appropriate for sleep recordings. The EEG cap was connected to a 4m long cable that allowed patients to leave the bed and move around the room. Four electrodes located near the eyes were used to monitor eye movements, and electrodes overlying the masseter muscles to record muscle tone^13^.

#### Preprocessing

EEG recordings underwent a thorough artefact removal procedure (Fig. S8). First, the EEG was bandpass filtered between 0.5 and 35 Hz. Then, the EEG corresponding to parasomnia episodes was extracted, as well as the EEG of the five minutes of sleep preceding movement onset. To remove additional artifacts, artifact subspace reconstruction (ASR)^58^ was applied to the EEG corresponding to the parasomnia episode (after movement onset). ASR is an advantageous method due to its capability to efficiently and automatically eliminate non-stationary artifacts of significant magnitude, irrespective of their distribution on the scalp or their consistency across the dataset^58–62^ and was recently validated for sleep ^63^. It works as an adaptive spatial filtering algorithm, which uses clean sections of EEG signal to produce a statistical model to reject artifactual EEG segments based on how much they deviate from the calibration signal in the principal component analysis subspace^58^. The artifact-free sections were automatically derived by the algorithm, based on standard deviations from a robust estimate of the EEG power distribution in each channel. A moving window of 768ms (1.5 × the number of channels) and a 25 standard deviation cutoff for rejection were used to avoid rejection of sleep features (such as slow waves persisting during the awakening/episode). The threshold was determined based on visual inspection to maximally remove noise and completely preserve sleep slow waves^58^. EEG analyses at the scalp level were performed on the innermost 175 channels to further limit artifact contamination. Artifactual channels were visually inspected, marked and interpolated with data from other channels using spherical splines (NetStation, Electrical Geodesic Inc). To exclude ECG and other artifacts, including ocular, movement and electrodermal activity, independent component analysis (ICA) was performed using EEGLAB routines on sleep and parasomnia episodes separately^64,65^.

#### Spectral power decomposition

The spectral power decomposition of the EEG signal was computed on a 40s timeframe centered on movement onset^13^, for non-overlapping 2s and 0.5s epochs in the delta (1-4 Hz) and beta (26-34 Hz) frequency bands, respectively. The frequency bands were chosen as representative for low-frequency power (slow wave activity) and high-frequency power respectively, based on our previous study on dreaming^33^, and to simplify presentation of results. Analyses of spectral power were based on the 20s of sleep preceding movement onset (“sleep EEG”, in light of our previous studies on the EEG correlates of dreaming^13^), and on the 20s window after movement onset (“parasomnia EEG”, for symmetry and because this timeframe approximately corresponded to the median duration of parasomnia episodes). The first four seconds after movement onset were excluded from analysis because of the frequent presence of residual movement artifacts in the EEG. Thus, the analysis for the parasomnia episode was restricted to the timeframe ranging from +4 to +20s after movement onset.

#### Source modelling

Source localization was performed using the GeoSource software (Electrical Geodesics, Inc., Eugene, Oregon), on filtered and preprocessed EEG segments. Individualized geocoordinates of the electrodes were used to construct the forward model. The source space was restricted to 2,447 dipoles distributed over 7×7×7-mm cortical voxels. The inverse matrix was computed using the standardized low-resolution brain electromagnetic tomography (sLORETA) constraint^66^. A Tikhonov regularization (λ=10^-2^) procedure was applied to account for the variability in the signal-to-noise ratio^66^.

#### Slow wave analysis

A previously validated slow wave detection algorithm^12,62^ was applied to the EEG signal. The EEG signal was baseline corrected, subtracting to each electrode signal its mean. Then it was re-referenced to the average of the two mastoids electrodes, downsampled to 128 Hz and bandpass filtered (0.5-4Hz; stop-band at 0.1 and 10 Hz) using a Chebyshev Type II filter (Matlab, Mathworks). The slow wave detection was performed for each channel separately and consisted in identificating the zero crossings, the negative deflections between two zero crossings were defined half-waves. Only half-waves with a duration between 0.25s and 1s were considered, no amplitude threshold was applied. After applying a moving average filter of 50ms, the determination of negative and positive peaks was based on the zero crossings of the derivative of the signal. Then for each slow wave different parameters were extracted: maximum negative peak amplitude, slope 1 (between the first zero crossing and the negative peak), slope 2 (between the negative peak and the second zero crossing), the number of negative peaks and the duration (time from the first and second crossing). Finally, the parameters of the waves contained by each instance were averaged and the density (expressed in number of slow waves per minute) was extracted.

#### Statistical Analyses

Statistical analyses were performed in R-Studio (version 1.1.463). Separate generalized linear mixed models were used to predict the presence of CE vs. NE or CEWR as a function of different variables, including the provoked or spontaneous nature of the episodes, the occurrence during the baseline or recovery sleep, the time passed since lights off, and the duration of the episodes. A generalized linear mixed model was also used to predict the spontaneous vs. provoked nature of the episode as a function of manifestations of surprise. For this model, a surprise index, ranging from 0 to 4 was calculated for each episode, with each of the following manifestations contributing one point to the score when it was present: sudden onset, startle-like movements, orienting behaviour, perplexity. Subject identity was always included as a random factor in all models.

To investigate how EEG activity predicted different categories of report (CE, CEWR, NE, etc.), generalized linear mixed models were computed for each electrode (scalp) or voxel (source) and timeframe (‘sleep’ and ‘parasomnia’). The frequency bands (delta and beta) were included as fixed factors, and subject identity and the nature of the awakening (spontaneous/provoked) as random factors. The Wald statistics (the squared ratio of the fixed factor estimate over its standard error) were obtained for each model, with an alpha level of 0.05. We opted for the one-sided statistics because we had a priori hypothesis, based on our previous work about the direction of results. In this sense the p-values resulting from the model were therefore divided by two. A cluster and probability-based correction was applied as follows: a dummy population was created by shuffling the labels between report categories 1000 times. For each permutation, the model was applied and neighbouring electrodes with p-values < 0.05 (according to one or two-tailed statistics) were identified as a cluster. Electrodes/voxels clusters were defined spatially, and neighbors were defined through a function from the Matlab toolbox fieldtrip using the triangulation-method, which calculates a triangulation based on a two-dimenstional projection of the sensor position. A cluster statistic was produced by averaging Wald statistics within each significant cluster and the maximal absolute cluster statistics was retained for each permutation. The real cluster statistics obtained in the real dataset were compared with the threshold of significance, which was set at the 97.5^th^ or 95^th^ percentile (according to one or two-tailed statistics) of the dummy cluster statistics distribution. The generalized linear mixed model formulas and the voxel/channels statistics are indicated in Tables S1 and S3, respectively.

## Funding and acknowledgements

This work was supported by grants of the Swiss National Science Foundation (Ambizione Grant PZ00P3_173955), the Théodore-Ott Foundation and the Foundation for the Advancement of Neurology awarded to F.S. The authors thank Jean-Baptiste Maranci and Henry Hebron for insightful discussions.

## Data availability

The EEG datasets generated and analysed in the current study can be requested from the corresponding author after the main publications resulting from this dataset have been completed and if the participants’ privacy is protected in accord with applicable Swiss laws and regulations.

## Code availability

The custom-made codes in Matlab, Mathworks (version 2015a) and R-Studio (version 1.1.463) are available upon request.

