## Supplementary information for "Non-REM parasomnia experiences share EEG correlates with dreams"

**Figure S1**

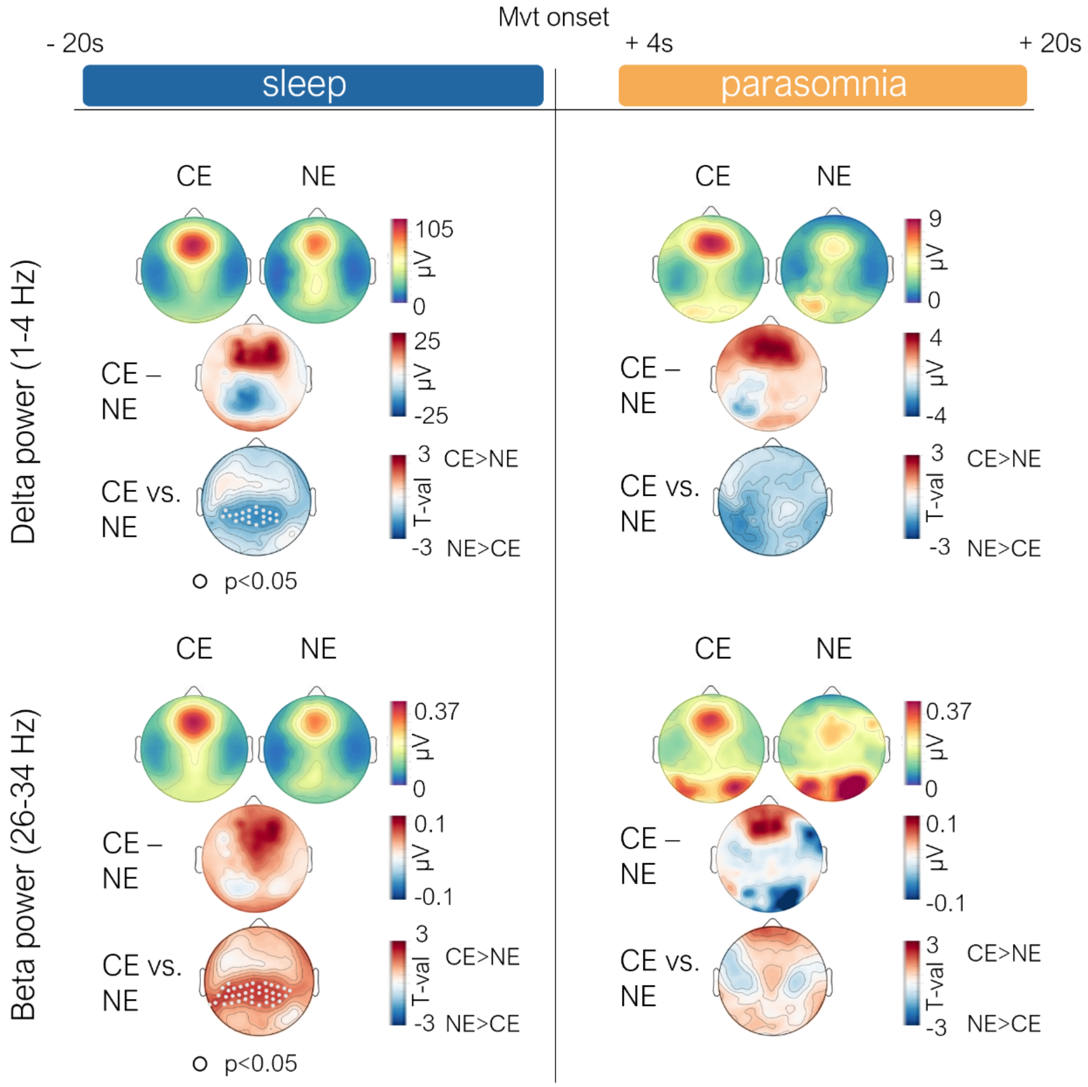


**Fig. S1: Conscious experience (CE) vs. no experience (NE): spectral power scalp level. A.** Left column**:** topographical distribution of absolute delta power (1-4 Hz) averaged across subjects for CE (n = 14) and NE (n = 7), of the absolute difference (CE minus NE), and of t-values (Wald statistics, CE (n = 32) vs. NE (n = 11)) at the scalp level for the 20 seconds of sleep preceding movement onset. Right; same as left for the period corresponding to the parasomnia episode, from 4 to 20 seconds after movement onset (CE (n = 31) vs. NE (n = 10)). 175 innermost channels are displayed. **B.** Same as A for beta power (26 – 34 Hz). Mvt onset= movement onset.

**Figure S2**

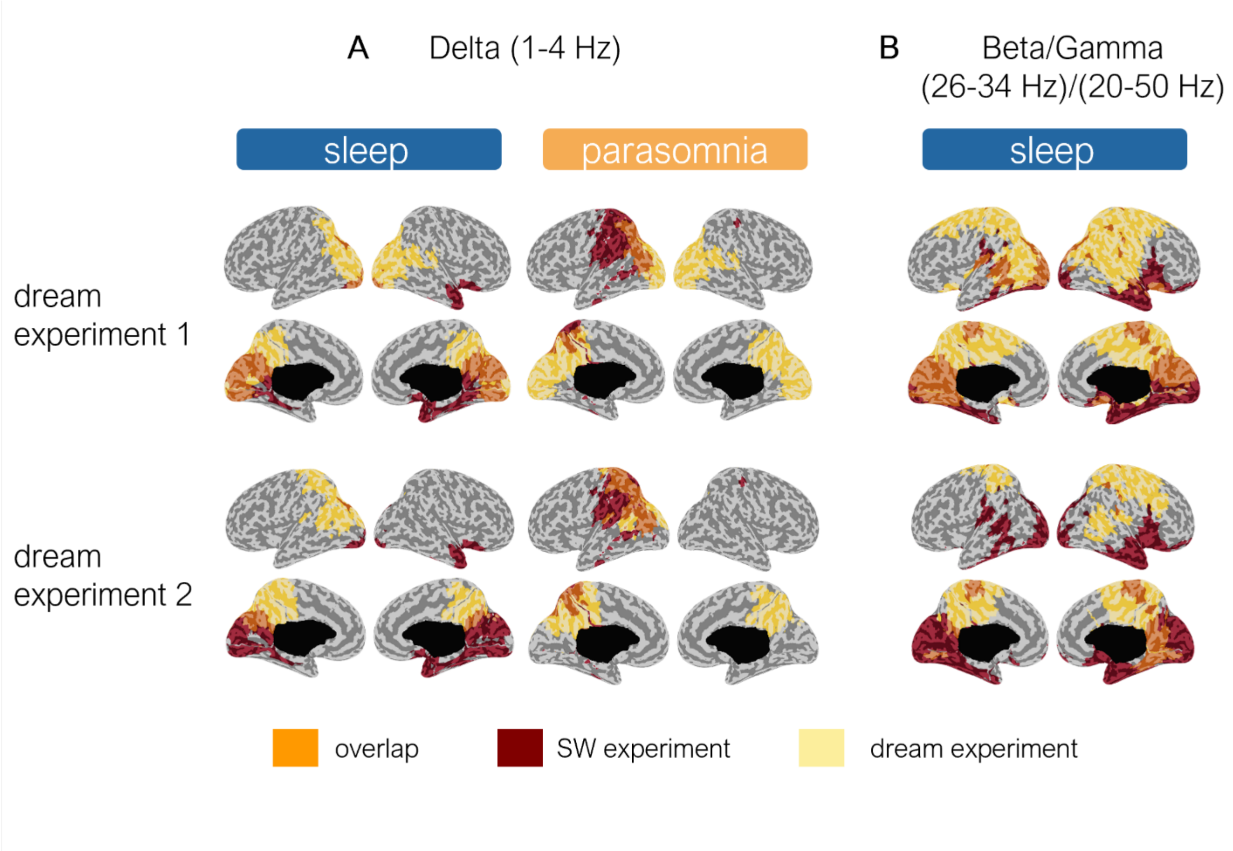


**Fig. S2:Conjunction maps: differences and overlap between** contrast conscious experience/no experience in the current study and the contrast dream experience/no experience in two previous experiments on the neural correlates of dreaming in healthy participants (Siclari et al., Nat Neurosci 2017) for low frequency power (A) and high-frequency power (B). Power in the high-frequencies was analyzed in the beta band in the current study (26-34 Hz), and in the gamma band (20-50Hz) in the previous study on dreaming. Statistics were not computed in the same manner (generalized linear mixed models in the current study, paired t-tests in the dream study).

#### Figure S3

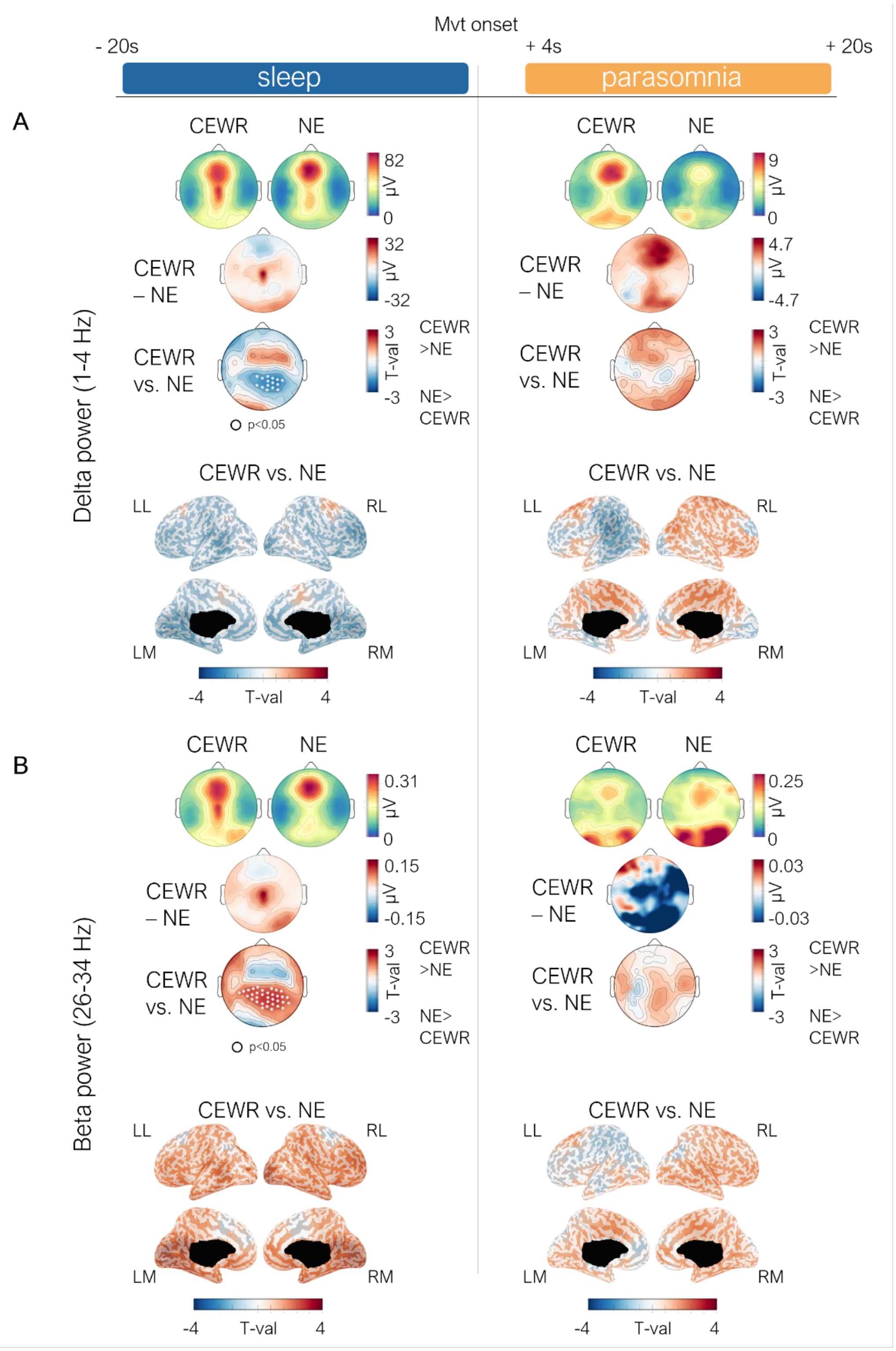


**Fig. S3. Conscious experience without recall of content (CEWR) vs. no experience (NE): spectral power.**
Left column: topographical distribution of absolute sleep delta power (1-4 Hz) averaged across subjects for CEWR (n = 8) and NE (n = 7), of the absolute difference (CEWR minus NE), and of t-values (Wald statistics, CEWR vs. NE) at the scalp and source level for 20 seconds of sleep preceding movement onset. Right; same as left for the period corresponding to the parasomnia episode from 4 to 20 seconds after movement onset. 175 innermost channels are displayed at the scalp level. LL, left lateral; RL, right lateral; LM, left medial; RM, right medial. B. Same as A for beta power (26–34 Hz). Mvt onset= movement onset.

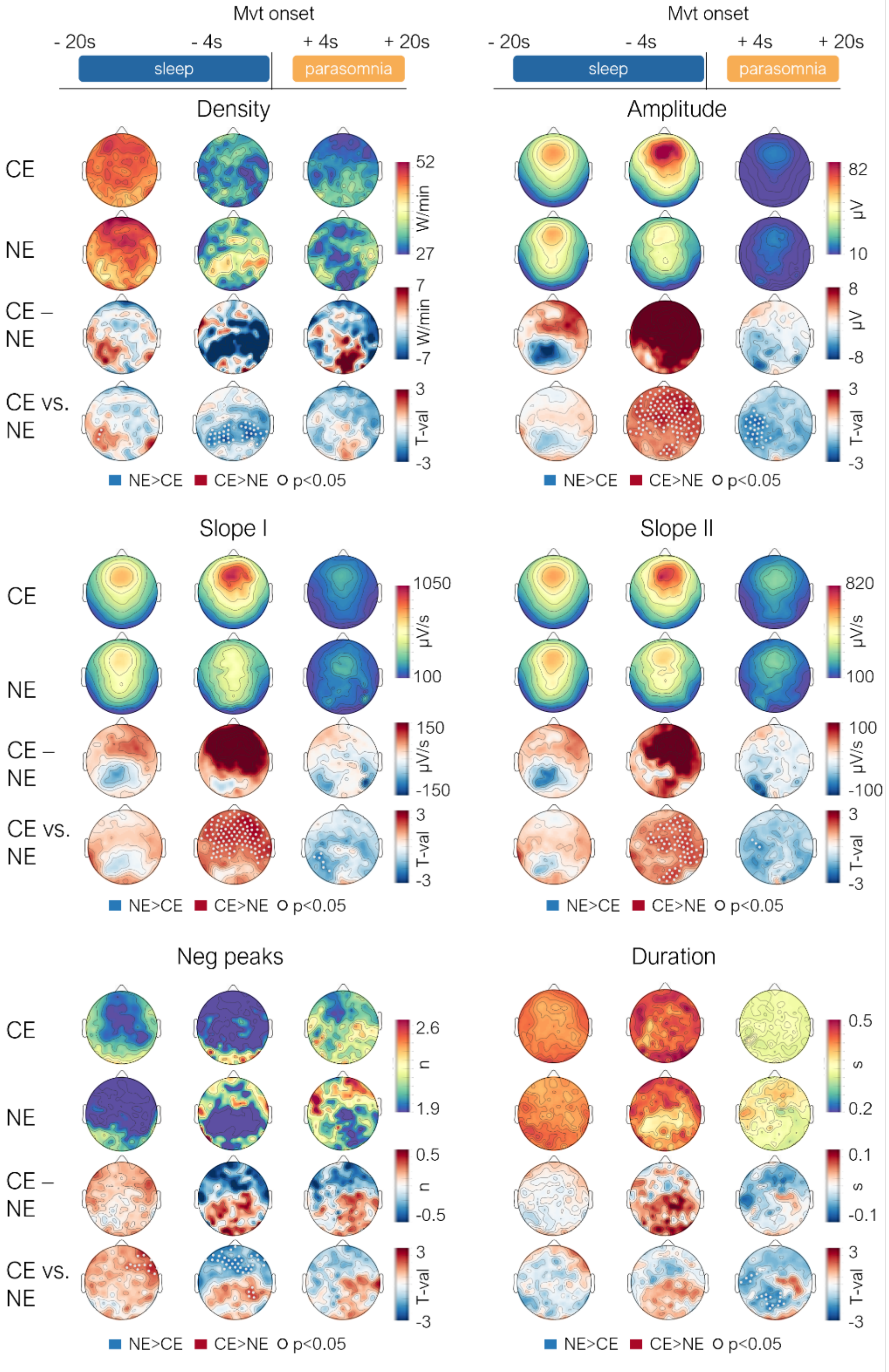


### Figure S4

**Fig.S4: Conscious experience (CE) vs. no experience (NE): slow wave parameters.** Topographical distribution of slow wave parameters averaged across subjects for CE (N=14 subjects, 32 episodes) and NE (n=7, 11 episodes), of their absolute difference (CE minus NE), and of t-values (Wald statistics, CE vs. NE) for three timeframes (sleep from -20 to -4s before movement onset (n= 43 trials), sleep from -4s to movement onset (n= 43 trials) and the parasomnia episode from +4s to +20s (n= 41 trials)). 175 innermost channels are displayed. Density=number of slow waves per minute, Amplitude= absolute amplitude of maximum negative slow wave peak. Slope I=positive to negative deflection of the slow wave, Slope II=slope of negative to positive deflection of slow wave, Neg Peaks= number of negative intrawave peaks, Duration = time between two zero-line crossings. Mvt onset= movement onset.

**Figure S5**

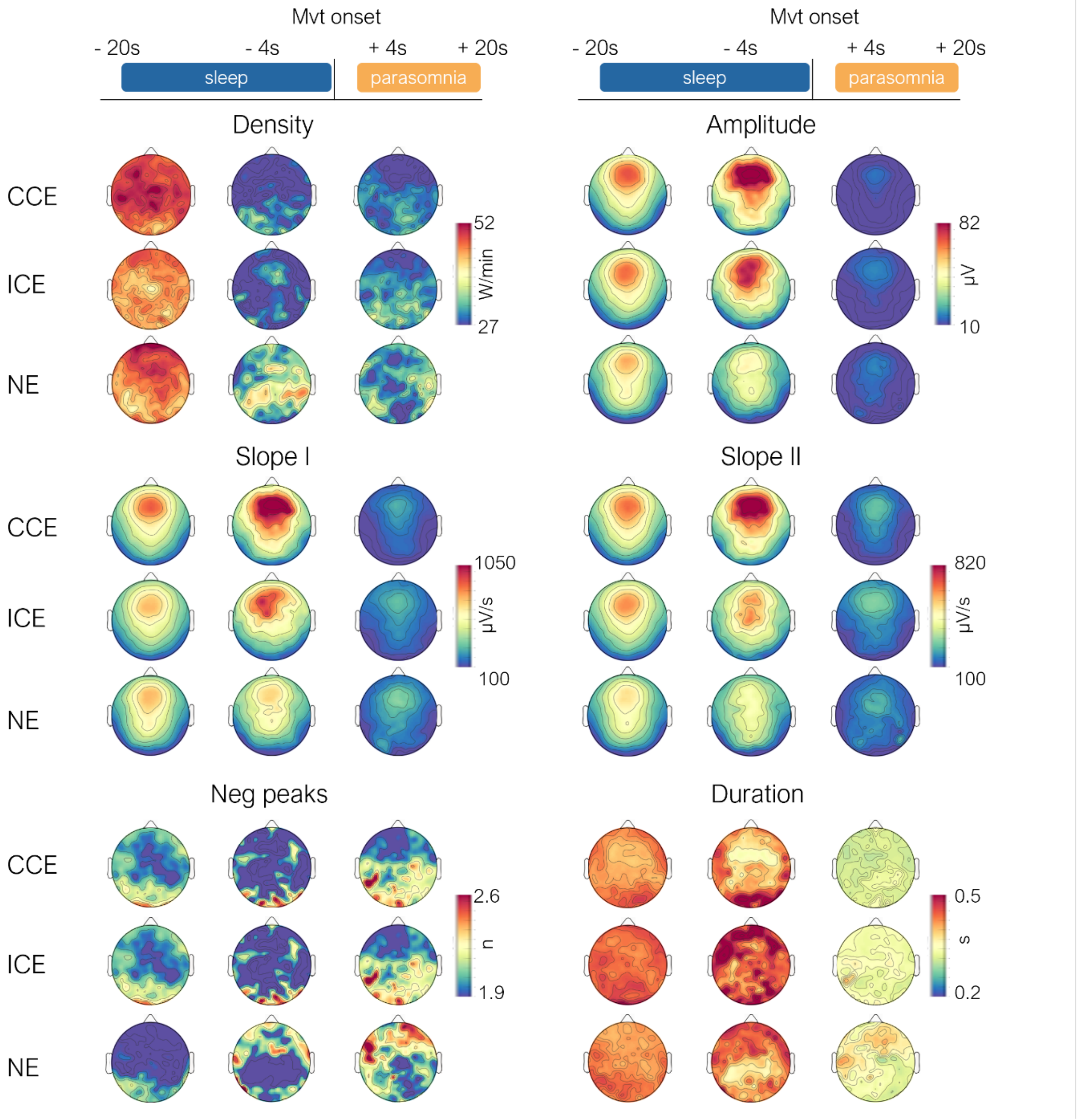


**Fig. S5: Coherent (CCE) and incoherent (ICE) conscious experiences vs. no experience (NE): slow wave parameters.** Topographical distribution of absolute slow wave parameters averaged across subjects for CCE (n=9 subjects, top row of each panel), ICE (n=9 subjects, middle row) and NE (n=7 subjects, bottom row), shown for three timeframes (sleep from -20 to -4s before movement onset, sleep from -4s to movement onset and the parasomnia episode from +4s to +20s). 175 innermost channels are displayed. Density=number of slow waves per minute, Amplitude= absolute amplitude of maximum negative slow wave peak. Slope I=positive to negative deflection of the slow wave, Slope II=slope of negative to positive deflection of slow wave, Neg Peaks= number of negative intrawave peaks, Duration = time between two zero-line crossings. Mvt onset= movement onset.

**Figure S6**

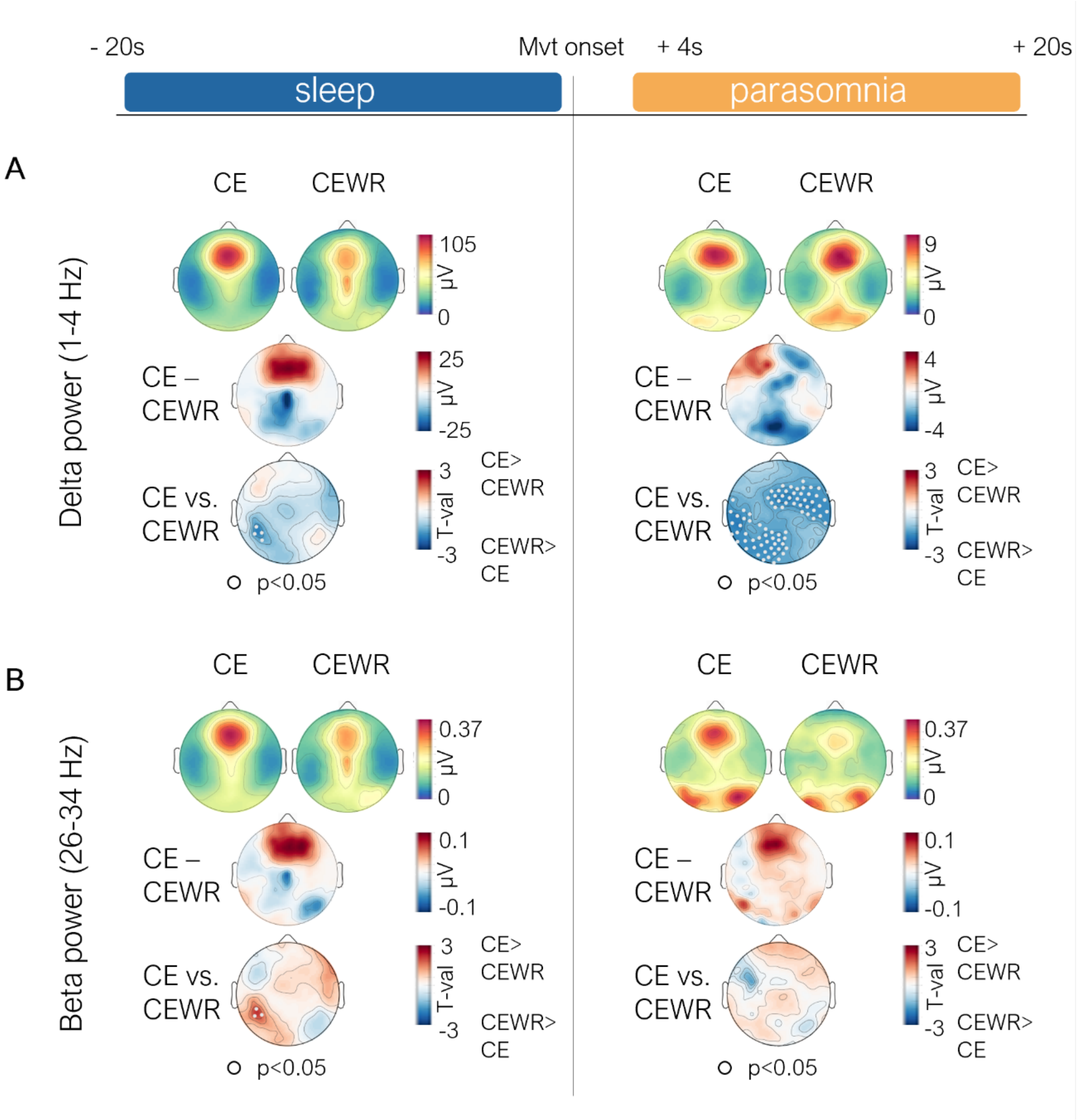


**Fig. S6. Conscious experience without recall of content (CEWR) vs. no experience (NE): spectral power.**
Left column: topographical distribution of absolute sleep delta power (1-4 Hz) averaged across subjects for CEWR (n = 8) and NE (n = 7), of the absolute difference (CEWR minus NE), and of t-values (Wald statistics, CEWR vs. NE) at the scalp and source level for 20 seconds of sleep preceding movement onset. Right; same as left for the period corresponding to the parasomnia episode from 4 to 20 seconds after movement onset. 175 innermost channels are displayed at the scalp level. LL, left lateral; RL, right lateral; LM, left medial; RM, right medial. B. Same as A for beta power (26–34 Hz).

**Figure S7**

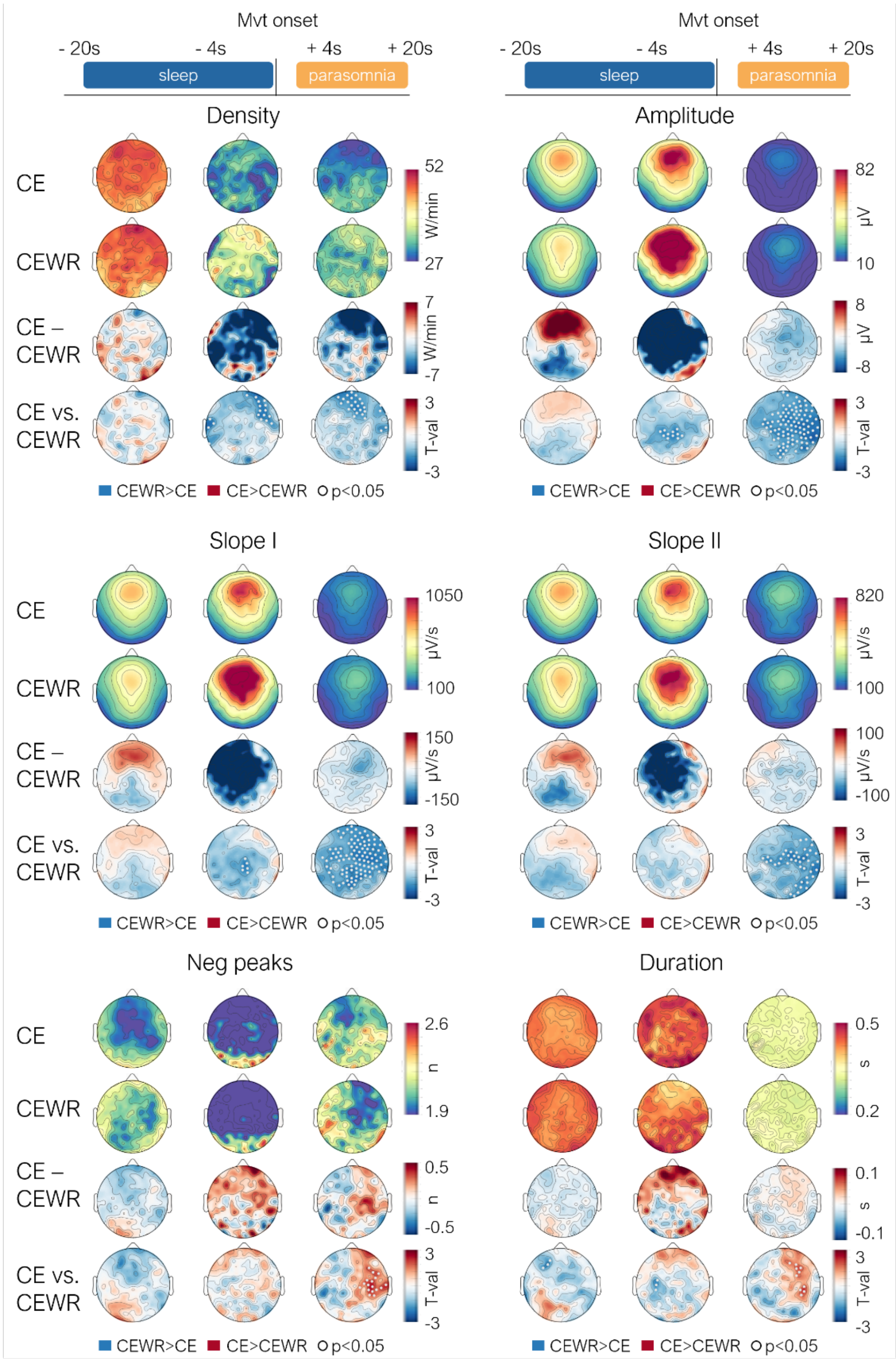


**Fig. S7: Conscious experience (CE) vs. no experience (NE): slow wave parameters.** Topographical distribution of slow wave parameters averaged across subjects for CE (N=14 subjects, 32 episodes) and CEWR (n=8, 17 episodes), of their absolute difference (CE minus CEWR), and of t-values (Wald statistics, CE vs. CEWR) for three timeframes (sleep from -20 to -4s before movement onset (n = 49 trails), sleep from -4s to movement onset (n = 49 trails)). 175 innermost channels are displayed. Density=number of slow waves per minute, Amplitude= absolute amplitude of maximum negative slow wave peak. Slope I=positive to negative deflection of the slow wave, Slope II=slope of negative to positive deflection of slow wave, Neg Peaks= number of negative intrawave peaks, Duration = time between two zero-line crossings. Mvt onset= movement onset.

**Figure S8**

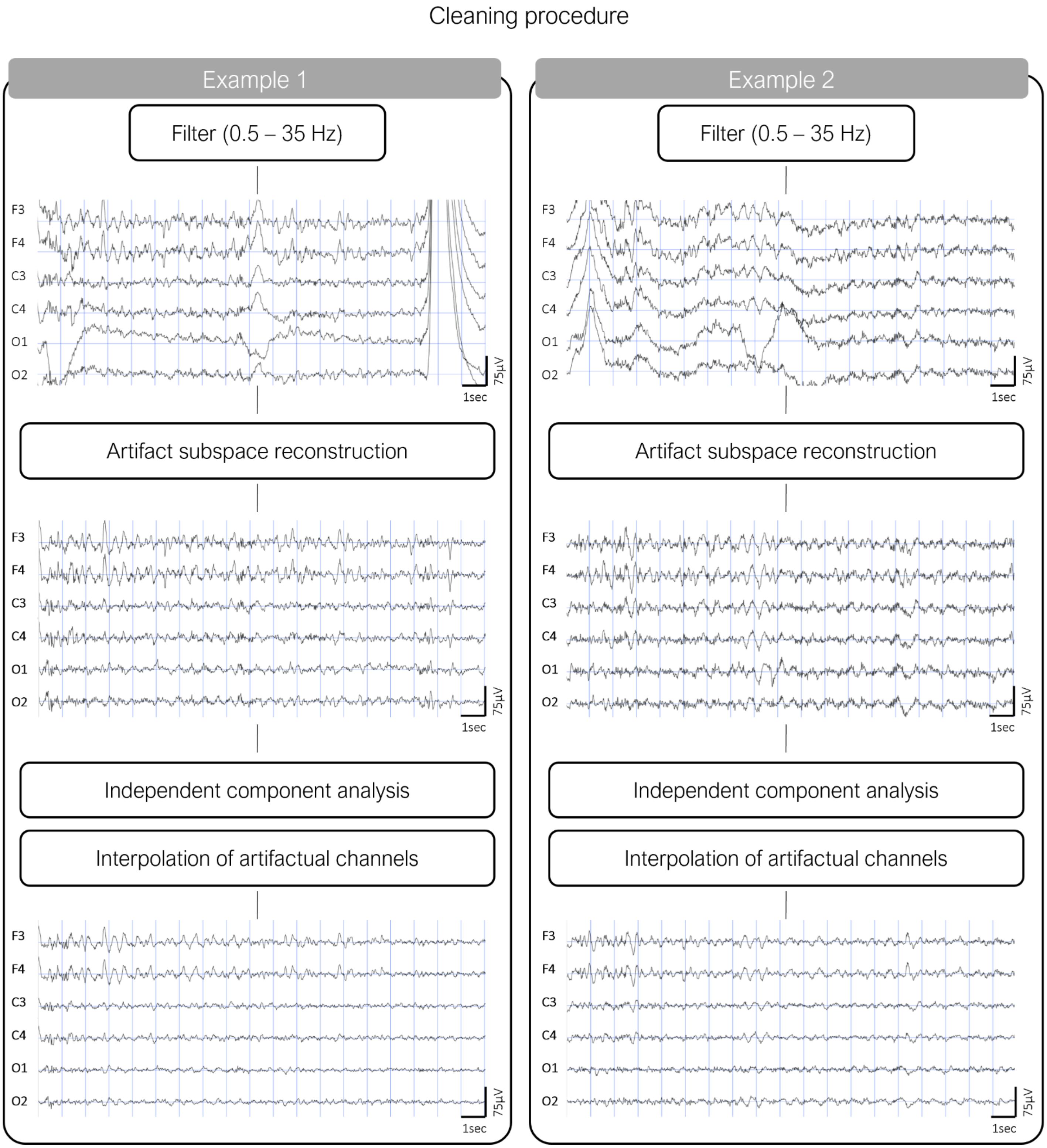


**Fig. S8: Artifact removal procedure:** Two examples of EEG traces ofNREM parasomnia episodes illustrating the main steps of the procedure. A selection of 6 channels (F3, F4, C3, C4, 01, 02) referenced to the linked mastoid is shown.

**Table S1**

| | Formula | Statistics | p |
|---|---|---|---|
| 1 | Prov/spont ~ surprise level + (1\|sub) | $X^2(1) = 0.219$ | 0.639 |
| 2 | CE/NE ~ prov/spont + (1\|sub) | $X^2(1) = 1.274$ | 0.264 |
| 3 | CE/CEWR ~ prov/spont + (1\|sub) | $X^2(1) < 0.01$ | 0.996 |
| 4 | CE/NE ~ BL/REC + (1\|sub) | $X^2(1) = 0.828$ | 0.362 |
| 5 | CE/CEWR ~ BL/REC + (1\|sub) | $X^2(1) = 1.277$ | 0.445 |
| 6 | CE/NE ~ time since lights off + (1\|sub) | $X^2(1) = 0.582$ | 0.784 |
| 7 | CE/CEWR ~ time since lights off + (1\|sub) | $X^2(1) = -0.997$ | 0.319 |
| 8 | CE/NE ~ time after first sleep onset + (1\|sub) | $X^2(1) = 0.333$ | 0.739 |
| 9 | CE/CEWR ~ time after first sleep onset + (1\|sub) | $X^2(1) = -0.935$ | 0.349 |
| 10 | CE/NE ~ duration + (1\|sub) | $X^2(1) = 1.832$ Fig.1C | 0.067 |
| 11 | CE/CEWR ~ duration + (1\|sub) | $X^2(1) = 0.063$ Fig.1C | 0.546 |
| 12 | CE clear/NE ~ duration + (1\|sub) | $X^2(1) = 2.24$ | 0.024 |
| 13 | CE/NE ~ log (delta) + log (beta) + (1\|sub) + (1\|prov/spont) | Fig.2, Fig.S1 | |
| 14 | CE/NE ~ slow wave parameter + (1\|sub) + (1\|prov/spont) | Fig.3, Fig.S4 | |
| 15 | CCE/NE ~ log (delta) + log (beta) + (1\|sub) | Fig.4A | |
| 16 | ICE/NE ~ log (delta) + log (beta) + (1\|sub) | Fig.4A | |
| 17 | CCE/NE ~ slow wave parameter + (1\|sub) | Fig.4B | |
| 18 | ICE/NE ~ slow wave parameter + (1\|sub) | Fig.4B | |
| 19 | CE/CEWR ~ log (delta) + log (beta) + (1\|sub) + (1\|prov/spont) | Fig.5, Fig.S6 | |
| 20 | CE/CEWR ~ slow wave parameter + (1\|sub) + (1\|prov/spont) | Fig.6, Fig.S7 | |
| 21 | CEWR/NE ~ log (delta) + log (beta) + (1\|sub) + (1\|prov/spont) | Fig.S3 | |

**Table S1**: Generalized linear mixed models used in this work, with reference to the figure displaying the results (if present). When consisting in a single value, statistics are reported in the column ‘statistics’ and their p-value in the column ‘p’. For the other analysis at the channel/voxel level, see Table S3.

#### Table S2

| Patient | Episode type | Report of experience | Observed behavior | Coherence |
|---|---|---|---|---|
| P2 | S | I was telling you that I had a secret, that I knew how to fall asleep without making any sound, so as to trick the recording machines. | Turns to the wall and says “You cannot?”, then, while lying prone and leaning on her forearms, whispers unintelligibly for a long time, before looking around and saying "I need to stop talking" then smiles. | Yes |
| P2 | S | I had to save a cockroach, or ladybugs from dying, from gliding down the wall. | Calls for someone, touches the wall, looks into the space between the wall and the bed, as if something had fallen there, continues to search and call for help. | Yes |
| P2 | S | I wanted to take a shower. I heard someone talking and I asked whether this person was talking to me, that’s all. | Opens eyes, looks puzzled, looks to her feet, looks to the right and says "are you talking to me?", then says "No", shakes her head, lies back on pillow and closes eyes. | Yes |
| P4 | S | Cookies | Sits up, says something, points to something with the left hand, then talks very fast and unintelligibly, sighs (or laughs), pauses as if listened to someone, then talks again, as if in a conversation. | No |
| P12 | S | I was looking for my baby daughter, she wasn’t in the bed anymore, I thought she had fallen off the bed, I think I even screamed for help. | Suddenly sits up, touches wall with right hand, looks under the covers, then under the bed, and cries "Help!" twice. | Yes |
| P14 | S | A piece of furniture falling down | Lies on side, suddenly opens eyes with frightened expression, gasps and then lies back down, looks puzzled. | Yes |
| P17 | S | I was about to fall asleep. I saw something that made me startle. | Suddenly sits up in bed and looks around, seems puzzled. | Yes |
| P17 | S | I think I saw someone just in front of me and that made me startle. | Suddenly sits up in bed and looks around, seems puzzled. | Yes |
| P17 | S | Going to eat a pizza with brother. | Suddenly opens eyes looks around, seems puzzled. Scratches forehead close to electrodes. | No |
| P2 | P | I dreamt that there was a man, a nurse in the back of the room, on the right. He said that I should not pay attention to him. | Lifts head and says "Hm?" twice, looks to the far right of the room several times. | Yes |
| P2 | P | It is always about taxes, I haven’t slept these days, this will cost me a lot in terms of bills. | Opens eyes, looks to the right rapidly, lifts head and torso, touches her face and eyes, looks around puzzled. | No |
| P4 | P | Green pastures | Says ‘Hm?’, then ‘wait’, makes a hand gesture as if wanted to stop something, then talks unintelligibly. | No |
| P5 | P | I was in the train, seeing landscapes passing by. | Sits up, looks at cable box, readjusts cable above head. Makes smacking sounds with mouth. | No |
| P9 | P | Christmas. A candle on the tree, like a child would draw it. It was outside. I was doing something with the tree, then I turned around and startled when you talked to me. | After alarm sound opens eyes and looks around, then at the ceiling, then talks (appears to have an imaginary conversation). Startles when experimenter starts talking to her. | Yes |
| P14 | P | I thought I had lost the EEG net and cables | Suddenly opens eyes, seems puzzled and worried, says: I don’t have anything on me anymore? Do I? | Yes |

**Table S2**: examples of representative parasomnia experiences and associated behaviors. S=spontaneously occurring episode. P=provoked episode. Coherence= coherence between report and behavior.

#### Table S3

| **Duration** (s) | CE | CEWR | NE |
|---|---|---|---|
| Average | 31.5 | 24 | 18.18 |
| Std | 19.9 | 18.3 | 12.15 |
| Median | 28.3 | 17 | 17 |
| Min | 3 | 9 | 4 |
| Max | 108 | 76 | 40 |

**Table S3: Parasomnia episode duration.** Average, standard deviation (std), median, minimum (min) and maximum (max) values for episodes with report of conscious experience (CE), report of conscious experience without recall (CEWR) and report of no experience (NE). All values are expressed in seconds.

#### Table S4

| | scalp: CE/NE ~ power + (1\|sub) + (1\|prov_spont) | source: CE/NE ~ power + (1\|sub) + (1\|prov_spont) | scalp: CE/CEWR ~ power + (1\|sub) +(1\|prov_spont) | source: CE/CEWR ~ power + (1\|sub) +(1\|prov_spont) | scalp: CCE/NE ~ power + (1\|sub) | scalp: CEWR/NE ~ power + (1\|sub) +(1\|prov_spont) |
|---|---|---|---|---|---|---|
| Delta sleep α = 0.05 | Fig. S1A c1(n = 20): 1.84 | Fig. 2A c1(n = 353): 1.84 | Fig. S6A c1(n = 4): 1.88 | Fig. 5A c1(n = 48): 1.86 | Fig. 4A c1(n = 4): 1.81 | Fig. S3A c1(n = 4): 1.86 |
| Beta sleep α = 0.05 | Fig. S1B c1(n = 38): 1.96 | Fig. 2B c1(n = 775): 1.9 | Fig. S6B c1(n = 3): 2.04 | Fig. 5B c1(n = 101): 1.85 | Fig. 4B c1(n = 5): 1.9 | Fig. S3B c1(n = 34): 1.96, c2(n = 3): 2.01 |
| Delta epi α = 0.05 | NS | Fig. 2A c1(n = 301): 1.88 | Fig. S6A c1(n = 6): 1.84 c2(n = 39): 1.84 | Fig. 5A c1(n = 515): 1.93 | Fig. 4A c1(n = 33): 1.83 | NS |

**Table S4**: summary of the statistics for the significant clusters (electrodes/voxels) emerging from the generalized linear mixed models with power spectral density as fixed factor. Formula of the model is reported in the top row. For each time frame (sleep/episode) and cluster index (c1: cluster 1, c2: cluster 2 …), cluster size (n =) and T-Wald statistics (averaged across the voxels/electrodes of the cluster) are reported. NS = non-significant.

**Table S5**

| PSG-parameter | Average | Std |
|---|---|---|
| Sleep latency (to N1, in min) | 15.35 | 14.25 |
| Recording time (min) | 469.44 | 42.59 |
| Time after sleep onset (min) | 453.14 | 42.40 |
| Wake after sleep onset (WASO) (min) | 29.08 | 31.65 |
| Total sleep time (TST) (min) | 424.02 | 38.17 |
| Sleep efficiency (%) | 93.19 | 6.00 |
| REM latency (min) | 110.55 | 36.65 |
| N1 (min) | 26.10 | 15.30 |
| N2 (min) | 230.86 | 38.98 |
| N3 (min) | 82.00 | 28.23 |
| N1 (% of TST) | 6.16 | 3.67 |
| N2 (% of TST) | 54.45 | 8.41 |
| N3 (% of TST) | 18.95 | 6.54 |
| REM (% of TST) | 20.11 | 4.95 |
| Arousal index (n/h) | 13.11 | 5.43 |
| Apnea-hypopnea index (AHI) (n/h) | 4.27 | 4.91 |
| Periodic limb movements index (PLMI) (n/h) | 3.06 | 6.16 |

**Table S5**: Sleep parameters patients obtained from clinical polysomnography (PSG) in 22 patients.

**Text S1**

*Additional information about patients*

Reported age of onset of parasomnia episodes was 9.1 ± 5.6 yrs (3-22). Mean self-reported frequency of parasomnia episodes was distributed as follows: ~once a month: 2 patients (9%), 2-3 times a month: 6 patients (27%); ~once a week: 6 patients (27%); 2-3 times a week 5 patients (23%) and almost every night: 3 patients (14%). All 22 patients (100%) had a history of confusional arousals. In addition, 13 patients (59%) had a history of both sleep terrors and sleepwalking, 6 patients (27%) only of sleepwalking and 2 patients only of sleep terrors (9%). A family history of NREM parasomnias was present in 12 patients (54%). Neurological and psychiatric comorbidities included migraine (n=1), sleep paralysis (n=1) idiopathic hypersomnia (n=1), a history of attention deficit hyperactivity disorder in childhood (n=1) and a history of febrile convulsions in childhood (n=1). Two patients had a periodic leg movements of sleep (PLMS) index greater than 15/h (23/h and 16.4/h), and five patients had an AHI index greater than 5/h, of which two had an AHI greater than 15/h (16.7/h and 15.5/h).

**Video S1**

Spontaneous parasomnia episode with report of no experience.

**Video S2**

Spontaneous parasomnia episode with report of experience that was judged coherent with the behavior displayed. ‘I was looking for my baby daughter, she wasn’t in the bed anymore, I thought she had fallen off the bed, I think I even screamed for help’.

**Video S3**

Spontaneous parasomnia episode with report of experience that was judged incoherent with the behavior displayed ‘I was going to eat pizza with my brother’.
